# Discussion-based DEI education to help create inclusive and open BME research lab environments

**DOI:** 10.1101/2022.03.30.486429

**Authors:** Ritika Naiknavare, Katharina Maisel

## Abstract

Diversity in teams has been shown to enhance creativity and innovation, particularly in teams where all members felt a sense of belonging. Creating an inclusive environment in a lab setting that provides a sense of belonging to all is challenging. This is particularly true in a field like Biomedical Engineering/Bioengineering where diversity is multifaceted and includes people’s diverse personal/cultural backgrounds and also diverse scientific backgrounds. In a research lab, there is additional diversity in training, as most labs contain trainees at different levels (undergrad, grad, postdoc, high school). To aid in creating a sense of belonging in a research lab, we have devised a novel initiative based upon open group discussions on diversity, equity, and inclusion topics. The initiative included a first presentation/discussion by the PI to set the stage for defining diversity, equity, and inclusion, and provided examples of existing DEI issues within our field, such as lack of racial diversity in awards like the NIH New Innovator Award and among conference speakers. After the initial presentation, trainee-led, bi-weekly-structured discussions were maintained. First, discussions focused broadly on any DEI related topic to enhance general knowledge on DEI issues and provide discussion comfort among the group. Second, there was a period to deepen knowledge of a specific topic (in our example, microaggressions), which was followed up with a period of discussions on potential solutions (such as how to react when observing a microaggression and what to do in response to realizing one’s own microaggression). Students reported that our discussions were the only ones they have had in their training thus far, and felt that these discussions made them feel like they belonged in the lab, made the lab more inclusive, enhanced their awareness of how to create inclusive spaces, and taught them about a variety of DEI topics. Overall, our DEI discussions have fostered open conversations within our lab group about DEI-related issues and topics at the lab, department, university, and countrywide levels and has established a space where students feel safe to voice their opinions and ask questions. We hope to ultimately use these DEI discussions to create actionable steps for addressing topics, e.g., microaggressions, in different scenarios that can be applied by group members in their future careers.

## Challenge

Diversity in engineering is still far below the general population and racism and sexism play significant roles in reducing both interest and continued pursuit of engineering careers by diverse groups ^1, 2^. Research has demonstrated that teams made up of diverse individuals (culturally and otherwise) outperform homogeneous teams and diverse individuals often pursue more creative approaches to problem solving ^3-5^. However, current diversity numbers in engineering, particularly at the level of those obtaining graduate degrees and within academic environments, do not reflect the general population (**Table 1**)^1^. Even in fields like Bioengineering/Biomedical Engineering, where close to 40% of doctoral degrees are earned by women, only 20-25% of new Assistant Professors are female ^6^. These numbers are further exacerbated for under-represented minorities (URM) and URM women – less than 25% of the already too low percentages of Blacks and Hispanics earning undergraduate and graduate degrees are women ^6^. Therefore, there is a significant need to create research environments that promote and maintain diversity, and in engineering particularly, to increase recruitment and retention of women and URM. Research has shown that even when participation is broadened, underrepresented groups are more likely to leave STEM, indicating that retention is a significant problem in creating a diverse STEM workforce ^7^. Perceptions and societal integration are at least partly at fault for poor retention of underrepresented groups in science ^7^ and creating inclusive environments has been recognized to be one of the keys to increase retention of URM in STEM. To foster an inclusive environment, we have developed a discussion-based strategy that includes student-led short (20-30 min), informative, and interactive presentations that combine the current state of the topic/issue with thought exercises and discussions, which can be incorporated into research group or similar team meetings.

**Table 1:** Percentage of women and people of black/African American or Hispanic race and ethnicity in the general population, earning engineering undergraduate or doctoral degrees, or tenure-track faculty at 4-year colleges or universities ^1^.

|  | Women | Black or African American | Hispanic |
| --- | --- | --- | --- |
| **General population** | 51% | 12% | 16% |
| **Undergraduate degrees** | 21% | 4% | 10% |
| **Doctoral degrees** | 24% | 2% | 3% |
| **Tenure-track faculty** | 16% | 3% | 5% |

## Novel Initiative

### Rationale

While there are several strategies to improve Diversity, Equity, and Inclusion within a community, our approach was to integrate DEI-related discussions into every other weekly lab meeting. These conversations served as a gateway to explore and learn about DEI-related issues and their relation to the STEM field in an informal, yet educational environment. While DEI discussions remain a novel concept, this form of executing DEI initiatives has seen success. Whether it has been through collegiate classes or departmental seminars, data has shown that conversational series such as our DEI discussions have paramount effects in improving diversity and inclusion. This discussion-based format has been seen to increase the understanding of the minority experience in STEM^8^, significantly increase self-perception and implicit bias^9^, and has been shown to increase feelings of inclusion within underrepresented minority groups^10^. In addition to social changes, DEI discussions have also created tangible change, such as increasing faculty hiring of underrepresented minorities and, overall, creating a more diverse workforce^11^. In summary, research indicates that an informal, group-based environment, as opposed to more formal and individually-based training, is much better suited for learning about and improving diversity, equity, and inclusion within academia.

### Structure

Our group DEI discussions have taken place every other week during the time allotted for our weekly lab meetings. These efforts were kicked off with an ‘orientation’ presentation on justice, equity, diversity, and inclusion within the field of Bioengineering/Biomedical Engineering to set the stage for why there is a need to continue the conversation and education on DEI topics. This presentation should include examples of representation in awards, grant funding, and higher level positions within the field, such as for example the winners of the NIH New Innovator Award. After setting the stage with this initial presentation, the following discussions are primarily student-led, where each of the undergraduate and graduate students are responsible for directing the discussion on a rotating basis. The student leading the discussion that week is responsible for creating a short, informative presentation on the topic at hand.Presentations should include a description of the problem is, who is affected by it, and questions to pose to the group to enhance discussion, self reflection, and develop mitigating strategies.

The content of such DEI presentations can vary from semester to semester and are chosen at the beginning of the term by the full lab group. Because of the reduced number of students and more focus on research, summer topics were typically broader and have helped to guide the topic of the following academic year (See **Table 2** for example topics). Each academic year, an overarching DEI topic is chosen; in past semesters, we have included topics such as microaggressions and emotional health. During the fall semester, discussions focus on understanding the topic and various viewpoints thereof, while the spring semester focuses on actionable items. Each presentation and discussion of the elected topic focuses on a different aspect of the topic to avoid repetition. For example, during Fall 2020 we covered microaggressions: we spent the first half of the semester weeks generally discussing the topic and understanding it through reading and discussing various journal and media articles, particularly focusing on how to identify microaggressions in various aspects of our life (**Table 2**). During Spring 2021, we focused on how to prevent and handle microaggressions in our daily lives. This process of first understanding and defining the topic, followed by identifying and learning how to combat it, is a strategy we plan on continuing to use to validate its effectiveness and recommend other interested groups to follow.

**Table 2:**
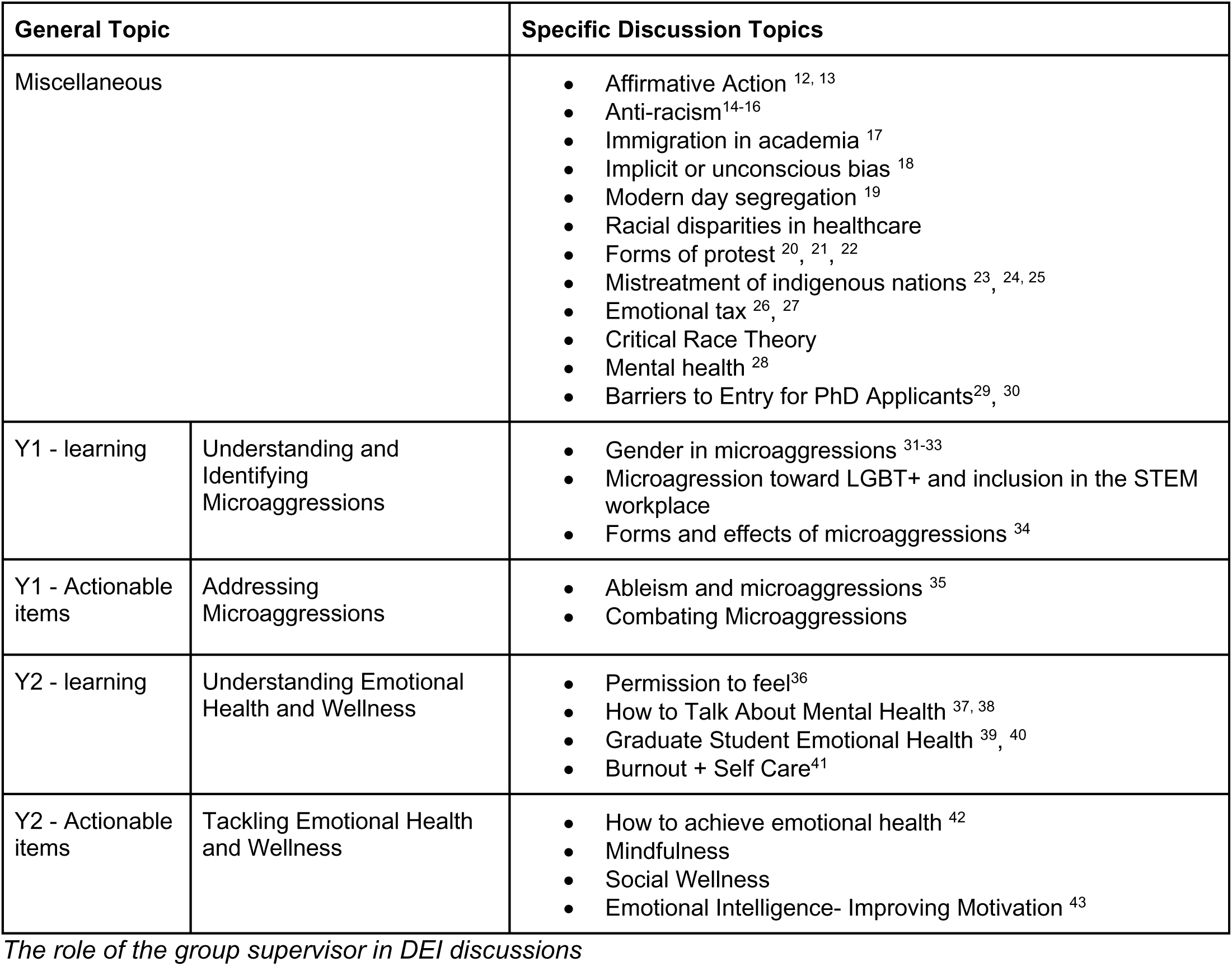
Example DEI Topics and resources.

| General Topic |  | Specific Discussion Topics |
| --- | --- | --- |
| Miscellaneous |  | Affirmative Action <sup>12, 13</sup><br>Anti-racism<sup>14-16</sup><br>Immigration in academia <sup>17</sup><br>Implicit or unconscious bias <sup>18</sup><br>Modern day segregation <sup>19</sup><br>Racial disparities in healthcare<br>Forms of protest <sup>20, 21, 22</sup><br>Mistreatment of indigenous nations <sup>23, 24, 25</sup><br>Emotional tax <sup>26, 27</sup><br>Critical Race Theory<br>Mental health <sup>28</sup><br>Barriers to Entry for PhD Applicants<sup>29, 30</sup> |
| Y1 - learning | Understanding and Identifying Microaggressions | Gender in microaggressions <sup>31-33</sup><br>Microaggression toward LGBT+ and inclusion in the STEM workplace<br>Forms and effects of microaggressions <sup>34</sup> |
| Y1 - Actionable items | Addressing Microaggressions | Ableism and microaggressions <sup>35</sup><br>Combating Microaggressions |
| Y2 - learning | Understanding Emotional Health and Wellness | Permission to feel<sup>36</sup><br>How to Talk About Mental Health <sup>37, 38</sup><br>Graduate Student Emotional Health <sup>39, 40</sup><br>Burnout + Self Care<sup>41</sup> |
| Y2 - Actionable items | Tackling Emotional Health and Wellness | How to achieve emotional health <sup>42</sup><br>Mindfulness<br>Social Wellness<br>Emotional Intelligence- Improving Motivation <sup>43</sup> |

### The role of the group supervisor in DEI discussions

DEI discussions need to be treated with sensitivity and thus it is crucial for the supervisor of the group to set the tone and establish a safe space. We have implemented a confidentiality agreement, where students know that whatever is shared within DEI discussions does not leave the room. Furthermore, supervisors should emphasize mutual respect within discussions - no labeling or judging of experiences should be accepted. Ideally, at the beginning of the semester supervisors should re-introduce the idea of confidentiality and mutual respect to maintain consistency. Supervisors are responsible for reminding group members that every experience is unique and all experiences have value. It is also crucial for the supervisor to intervene if inappropriate comments are made. While it is important to encourage questions and sharing of experiences, it is also vital not to push group members beyond their own comfort level. Group members may have had negative experiences with some of these sensitive topics and it is important to empower group members without any feeling of pressure. Additionally, it can be helpful to show vulnerability (but not overly expressing emotions) through sharing personal experiences on related topics. For example, discussions often start with silence from the group and sharing of personal experiences of the supervisor was usually able to break the ice. Sharing and reflecting on experiences of the supervisor can also help group members recognize that they may not be alone in their struggles, and difficult experiences do not have to prevent them from achieving their career goals. Finally, it is vital to show that the group values all contributions and topics brought up, which can be done by, for example, thanking group members for leading the discussion and sharing their experiences.

## Reflection

### Results from survey on DEI discussions

To better understand the effectiveness of our DEI discussions, we surveyed past and present lab members who have participated in discussions using an anonymous online survey, approved by the University of Maryland Institutional Review Board. Ten students participated in the survey (**Table 3**) that included basic demographics such as sex and race, which students could self-identify as. Seven participants self-identified as women, one participant chose not to say, and 2 self-identified as men (note: participants were given additional options including non-binary, gender-queer, transgender, other, and prefer not to say). Participants were asked to self-identify as white or person of color (or other), and had the option to identify as part of a traditionally underrepresented group in STEM. We want to note that participants who identified as persons of color largely also identified as Asian or Asian American and two participants chose not to disclose/prefer not to say. Participants were then asked to rate agreement with 12 statements from 1 – strongly disagree to 5 – strongly agree. Participants largely viewed the discussions positively (**Table 4**), with scores ranging mostly from 3-5. Students rated that they agree they learned a lot, felt comfortable sharing their thoughts, and felt discussions were valuable and respectful (**Table 4**). Interestingly, non-URM participants appeared to find discussions more valuable and increased sense of belonging compared to URM participants, though both groups rated both statements between neutral and strongly agree, on average (**Table 4**). On average, URM participants reviewed the discussions slightly less favorably than non-URM participants, as indicated by the significantly higher average of all questions. However, both URM and non-URM participant averages were between agree-strongly agree range, indicating that all participants found discussions valuable. Participants who identified as women also rated questions about comfort levels within discussions (e.g., “I feel comfortable sharing my thoughts”, “I feel respected during the discussions”, “I feel like my views and opinions were heard”) slightly more positively than men, though the number of male participants limited our analysis (**Table 5**). Additionally, participants identifying as white rated most questions more favorably than participants who identified as persons of color or other, as indicated by the significantly higher average of all questions for white persons, suggesting that there is still room for improvement. Overall, our initial data from survey participants indicate that DEI discussions may enhance participant feeling of belonging and value, suggesting that these discussions could contribute to a more inclusive environment within a research or other similar working group.

**Table 3:**
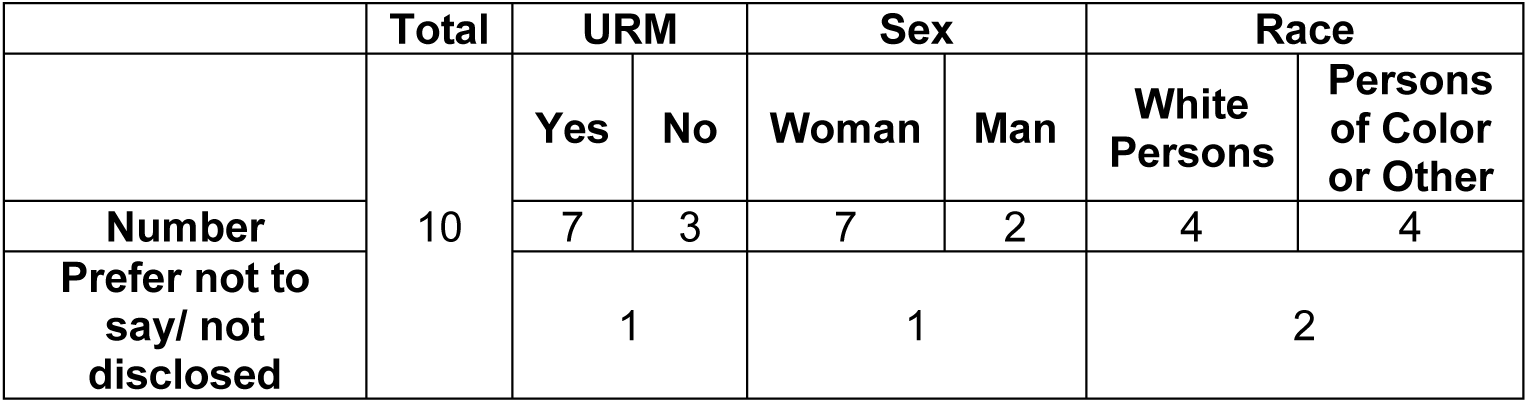
Summary of participant demographics.

|  | Total | URM |  | Sex |  | Race |  |
| --- | --- | --- | --- | --- | --- | --- | --- |
|  |  | Yes | No | Woman | Man | White Persons | Persons of Color or Other |
| Number | 10 | 7 | 3 | 7 | 2 | 4 | 4 |
| Prefer not to say/ not disclosed |  | 1 |  | 1 |  | 2 |  |

**Table 4:** Summary of survey on the effectiveness of discussions on diversity, equity, and inclusion in people who identify as underrepresented minority (URM) or not (non-URM). Participants were asked to rank the statements from 1-5, where 1 is strongly disagree, 2 somewhat disagree, 3 neutral, 4 somewhat agree, and 5 strongly agree. * indicates p < 0.05 comparing averages from URM and non-URM analyzed using student’s t-test.

| During the bi-weekly diversity, equity, and inclusion discussions: | Overall | URM | Non-URM |
| --- | --- | --- | --- |
| I learn a lot. | 3.7 ± 1.2 | 3.7 ± 1.4 | 3.7 ± 0.6 |
| I feel comfortable sharing my thoughts. | 3.8 ± 1.1 | 3.7 ± 1.4 | 4 ± 0 |
| I feel like my views and opinions were heard. | 4.3 ± 1.3 | 4.1 ± 1.5 | 4.7 ± 0.6 |
| I feel the discussions added to my feeling of belonging. | 3.7 ± 1.3 | 3.4 ± 1.4 | 4.3 ± 0.6 |
| I find these discussions valuable. | 4.1 ± 1.4 | 3.9 ± 1.7 | 4.7 ± 0.6 |
| I feel respected during the discussions. | 4.4 ± 1.3 | 4.3 ± 1.5 | 4.7 ± 0.6 |
| I would recommend this discussion format to other teams (labs and otherwise). | 4.2 ± 1.4 | 4.1 ± 1.6 | 4.3 ± 1.2 |
| I feel that these discussions contributed to my ability to relate to my labmates. | 4 ± 1.2 | 4 ± 1.4 | 4 ± 1 |
| These discussions make me feel that diversity, equity, and inclusion are valued in the lab working environment. | 4.7 ± 0.9 | 4.6 ± 1.1 | 5 ± 0 |
| I feel like I can bring up topics without judgment from others. | 4.1 ± 1.3 | 3.9 ± 1.5 | 4.7 ± 0.6 |
| I feel like I can make mistakes in these discussions. | 3.9 ± 1 | 3.7 ± 1.1 | 4.3 ± 0.6 |
| These discussions make me feel like I can bring my whole self to the lab working environment. | 4 ± 0.8 | 4 ± 1 | 4 ± 0 |
| Average from all statements | 4.1 ± 0.3 | 4 ± 0.3* | 4.4 ± 0.4 |

**Table 5:** Summary of survey on the effectiveness of discussions on diversity, equity, and inclusion broken down by demographic and sex. Participants were asked to rank the statements from 1-5, where 1 is strongly disagree, 2 somewhat disagree, 3 neutral, 4 somewhat agree, and 5 strongly agree. * indicates p < 0.05 comparing averages from White Persons and Persons of Color or Other analyzed using student’s t-test.

| During the bi-weekly diversity, equity, and inclusion discussions: | Overall | Female persons | Male persons | White Persons | Persons of Color or Other |
| --- | --- | --- | --- | --- | --- |
| I learn a lot. | 3.7 ± 1.2 | 4 ± 1.2 | 3.5 ± 0.7 | 4.3 ± 0.5 | 4 ± 1.2 |
| I feel comfortable sharing my thoughts. | 3.8 ± 1.1 | 4.14 ± 0.7 | 4 ± 0 | 4.3 ± 0.5 | 4.3 ± 0.5 |
| I feel like my views and opinions were heard. | 4.3 ± 1.3 | 4.71 ± 0.5 | 4.5 ± 0.7 | 4.8 ± 0.5 | 4.5 ± 0.6 |
| I feel the discussions added to my feeling of belonging. | 3.7 ± 1.3 | 4 ± 1 | 4 ± 0 | 4.5 ± 0.6 | 4 ± 0 |
| I find these discussions valuable. | 4.1 ± 1.4 | 4.3 ± 1.5 | 4.5 ± 0.7 | 5 ± 0 | 4.5 ± 0.6 |
| I feel respected during the discussions. | 4.4 ± 1.3 | 4.9 ± 0.4 | 4.5 ± 0.7 | 5 ± 0 | 4.5 ± 0.6 |
| I would recommend this discussion format to other teams (labs and otherwise). | 4.2 ± 1.4 | 4.7 ± 0.8 | 4 ± 1.4 | 5 ± 0 | 4.5 ± 1 |
| I feel that these discussions contributed to my ability to relate to my labmates. | 4 ± 1.2 | 4.3 ± 1.1 | 4 ± 1.4 | 4.8 ± 0.5 | 4.3 ± 1 |
| These discussions make me feel that diversity, equity, and inclusion are valued in the lab working environment. | 4.7 ± 0.9 | 5 ± 0 | 5 ± 0 | 5 ± 0 | 5 ± 0 |
| I feel like I can bring up topics without judgment from others. | 4.1 ± 1.3 | 4.4 ± 0.8 | 4.5 ± 0.7 | 4.5 ± 1 | 4.3 ± 0.5 |
| I feel like I can make mistakes in these discussions. | 3.9 ± 1 | 4 ± 0.8 | 4.5 ± 0.7 | 4.3 ± 1 | 3.8 ± 0.5 |
| These discussions make me feel like I can bring my whole self to the lab working environment. | 4 ± 0.8 | 4.3 ± 0.5 | 4 ± 0 | 4.3 ± 0.5 | 4.3 ± 0.5 |
| Average from all statements | 4.1 ± 0.3 | 4.4 ± 0.4 | 4.3 ± 0.4 | 4.6 ± 0.3* | 4.3 ± 0.3 |

### Future thoughts and improvements

We have found that DEI discussion can serve as a useful exercise to enhance student awareness about DEI topics and can contribute to an inclusive and positive working group environment. Despite the positive evaluation, we believe several items can help improve discussions for those interested in performing these with their own working groups or research teams. Since these discussions will happen regularly over a longer period of time, a more formal structure might be beneficial ^44^. After kicking off this effort in our group, we decided to have a two-phase strategy, spending one semester introducing the topic we have chosen for that year, and the second semester focusing on strategies to overcome barriers relating to the topic. In addition to having this broader structure, we have noticed that over time the discussion-based presentations can morph into more classic lecture-style presentations. To help prevent this, we plan to implement regular reminders on the structure of DEI discussions at the beginning of each semester. It may also be beneficial for the group leader or more-experienced group members to lead the first discussion, which can serve as an example for future discussions.

From our survey, we also found that several participants suggested more focus on actionable items would enhance the discussions and provide a platform to not only push for change within individual people, but also pursue change at departmental and institutional levels. One of the challenges with this suggestion is that while data on, for instance, representation or mental health, are widely available, it is more difficult to find – and implement – actionable items that can be done at an individual level. One suggestion we have to address this problem is to include brainstorming sessions within the discussion that focus on how individuals can contribute to change at broader levels. This may be particularly useful for topics where fewer obvious individual solutions exist. Additionally, it is important that discussions also include institutional or company resources aimed to address such challenges, whether these are in the form of additional trainings that are available or surveys/feedback forms that exist to provide trainees or colleagues with an opportunity to report incidents.

In summary, we have presented a structured, discussion-based DEI education tool that can aid in building an inclusive lab environment. These regular student-led discussions integrated into the context of official group meetings can help improve lab/research experiences, particularly for underrepresented groups, and create a more open lab environment. We aim to continue to develop this method for small and larger group settings.

## References

1. Statistics NCfSaE. Women, Minorities, and Persons with Disabilities in Science and Engineering. In: Foundation NS, editor. March 08, 20192019.

2. Erosheva EA, Grant S, Chen MC, Lindner MD, Nakamura RK, Lee CJ. NIH peer review: Criterion scores completely account for racial disparities in overall impact scores. Sci Adv. 2020;6(23):eaaz4868. Epub 2020/06/03. doi: 10.1126/sciadv.aaz4868. PubMed PMID: 32537494; PMCID: PMC7269672.

3. Hofstra B, Kulkarni VV, Munoz-Najar Galvez S, He B, Jurafsky D, McFarland DA. The Diversity-Innovation Paradox in Science. Proc Natl Acad Sci U S A. 2020;117(17):9284–91. Epub 2020/04/14. doi: 10.1073/pnas.1915378117. PubMed PMID: 32291335; PMCID: PMC7196824.

4. Watson WE, Johnson L, Zgourides GD. The influence of ethnic diversity on leadership, group process, and performance: an examination of learning teams. International Journal of Intercultural Relations. 2002;26(1):1 – 16. doi: https://doi.org/10.1016/S0147-1767(01)00032-3. PubMed PMID: WATSON20021.

5. Moon T. The effects of cultural intelligence on performance in multicultural teams. Journal of Applied Social Psychology. 2013;43(12):2414–25.

6. Institute of Education Sciences: National Center for Education Statistics. 2018. Available from: https://nces.ed.gov/ipeds/use-the-data.

7. Tennial RE, Solomon ED, Hammonds-Odie L, McDowell GS, Moore M, Roca AI, Marcette J. Formation of the Inclusive Environments and Metrics in Biology Education and Research (iEMBER) Network: Building a Culture of Diversity, Equity, and Inclusion. CBE Life Sci Educ. 2019;18(1):mr1. doi: 10.1187/cbe.18-03-0042. PubMed PMID: 30735086; PMCID: PMC6757218.

8. Reese AJ. An Undergraduate Elective Course That Introduces Topics of Diversity, Equity, and Inclusion into Discussions of Science. J Microbiol Biol Educ. 2020;21(1). Epub 20200410. doi: 10.1128/jmbe.v21i1.1947. PubMed PMID: 32313603; PMCID: PMC7148155.

9. Harrison-Bernard LM, Augustus-Wallace AC, Souza-Smith FM, Tsien F, Casey GP, Gunaldo TP. Knowledge gains in a professional development workshop on diversity, equity, inclusion, and implicit bias in academia. Adv Physiol Educ. 2020;44(3):286–94. doi: 10.1152/advan.00164.2019. PubMed PMID: 32484403; PMCID: PMC7642839.

10. Bernstein R, Bilimoria D. Diversity perspectives and minority nonprofit board member inclusion. Equality Diversity and Inclusion. 2013;32(7):636-+. doi: 10.1108/EDI-02-2012-0010. PubMed PMID: WOS:000213028000001.

11. Harpe J, Safdieh J, Broner S, Strong G, Robbins M. Creating a Neurology Department Diversity, Equity and Inclusion Committee. Neurology. 2021;96(15). PubMed PMID: WOS:000729283600141.

12. Gersen JS. The Uncomfortable Truth About Affirmative Action and Asian-Americans 2017 [updated 2017-08-10]. Available from: https://www.newyorker.com/news/news-desk/the-uncomfortable-truth-about-affirmative-action-and-asian-americans.

13. Menand L. The Changing Meaning of Affirmative Action. The New Yorker. 2020.

14. North A. What it means to be anti-racist. Vox. 2020.

15. How to be anti-racist: it’s more than books, quotes and Blackout Tuesday. Youtube: Washington Post; 2020.

16. 10 Ways To Promote Anti-Racism In The Workplace | Forbes. YouTube: Forbes; 2020.

17. Cole JR. Why American Universities Need Immigrants: @theatlantic; 2017 [updated 2017-03-07]. Available from: https://www.theatlantic.com/education/archive/2017/03/american-universities-need-immigrants/518814/.

18. Cover Story | Implicit Bias: Recognizing the Unconscious Barriers to Quality Care and Diversity in Medicine - American College of Cardiology2022.

19. Green EL. Within integrated schools, de facto segregation persists: @baltimoresun; 2022. Available from: https://www.baltimoresun.com/news/investigations/bs-md-school-segregation-series-howard-20170325-story.html.

20. Jackson KC. The Double Standard of the American Riot: @theatlantic; 2020 [updated 2020-06-01]. Available from: https://www.theatlantic.com/culture/archive/2020/06/riots-are-american-way-george-floyd-protests/612466/.

21. Lopez G. How violent protests against police brutality in the ‘60s and ‘90s changed public opinion: Vox.com; 2020 [updated 2020-06-02; cited 2022]. Available from: https://www.vox.com/2020/6/2/21275901/police-violence-riots-jacob-blake-kenosha-wisconsin.

22. Square ZP. When Is Rioting the Answer? : @TIME; 2015 [cited 2022]. Available from: https://time.com/3951282/riot-violence-use-american-history/.

23. Regan S. 5 Ways The Government Keeps Native Americans In Poverty: @forbes; 2022. Available from: https://www.forbes.com/sites/realspin/2014/03/13/5-ways-the-government-keeps-native-americans-in-poverty/.

24. Levin ST. ‘This is all stolen land’: Native Americans want more than California’s apology: @guardian; 2019 [updated 2019-06-21]. Available from: http://www.theguardian.com/us-news/2019/jun/20/california-native-americans-governor-apology-reparations.

25. NARF’s Five Priorities: Native American Rights Fund. Available from: https://www.narf.org/our-work/.

26. Catalyst. New Catalyst Report Reveals “Emotional Tax” Experienced by People of Colour in Corporate Canada: @PRNewswire; 2019 [cited 2022]. Available from: https://www.prnewswire.com/news-releases/new-catalyst-report-reveals-emotional-tax-experienced-by-people-of-colour-in-corporate-canada-300889953.html.

27. Florentine S. How to lower the Emotional Tax on minority employees 2018 [cited 2022]. Available from: https://www.cio.com/article/228507/how-to-lower-the-emotional-tax-on-minority-employees.html.

28. Loissel E. Mental Health in Academia: A question of support: eLife Sciences Publications Limited; 2019 [updated 2019-10-23]. Available from: https://elifesciences.org/articles/52881.

29. Wijesingha R. The Time Tax Put on Scientists of Color: Springer Nature Limited; 2020 [cited 2022].

30. Dolan EL. Institutional Interventions that Remove Barriers to Recruit and Retain Biomedical PhD Students. The American Society for Cell Biology. 2018;17(2). doi: 10.1187/cbe.17-09-0210.

31. Yang Y. Gendered Microaggressions in Science, Technology, Engineering, and Mathematics. In: Carroll DW, editor. Kansas State University: Kansas State University; 2022.

32. How Microagressions Are Like Mosquito Bites. YouTube: Fusion Comedy; 2016.

33. ADVANCE UoNH. Making the Invisible Visible: Gender Microaggressions. In: Hampshire UoN, editor.: University of New Hampshire.

34. Crandall J. “Am I overreacting?” Understanding and Combating Microaggressions -Higher Education Today: @ACEducation; 2016 [updated 2016-07-27; cited 2022]. Available from: https://www.higheredtoday.org/2016/07/27/understanding-and-combatting-microaggressions-in-postsecondary-education/.

35. Harris L. Exploring the Effect of Disability Microaggressions on Sense of Belonging and Participation in College Classrooms: Utah State University; 2017.

36. Brackett M. Permission to Feel: Unlocking the Power of Emotions to Help Our Kids, Ourselves, and Our Society Thrive: Celadon Books 2019.

37. Sword R. How to Talk About Mental Health: @hst; 2021 [updated 2021-02-19; cited 2022]. Available from: https://www.highspeedtraining.co.uk/hub/how-to-talk-about-mental-health/.

38. OK MI. Scenarios. makeitok.org; 2016.

39. Forrester N. Mental health of graduate students sorely overlooked. Nature. 2021;595(7865):135–7. doi: doi:10.1038/d41586-021-01751-z.

40. Puri P. The Emotional Toll of Graduate School. Scientific American. 2019.

41. Freudenberger HJ. Burnout: The High Cost of High Achievement: Bantam Books; 1981.

42. Wooll M. How to achieve emotional health2021.

43. Pettit M. 7 Powerful Ways to Increase Self-Motivation. Thrive. 2020.

44. Clark D, D’Angelo C, Menekse M. Initial Structuring of Online Discussions to Improve Learning and Argumentation: Incorporating Students’ Own Explanations as Seed Comments Versus an Augmented-Preset Approach to Seeding Discussions. Journal of Science Education and Technology. 2009;18(4):321–33. doi: 10.1007/s10956-009-9159-1. PubMed PMID: WOS:000270140300003.

